# Microglial Heterogeneity in a Mouse Model of Alzheimer’s Disease: A Mass Cytometry Analysis

**DOI:** 10.1101/404178

**Authors:** EM Fountain, TJ Esparza, O Malkova, ST Oh, DL Brody

**Author notes:** To whom correspondence should be addressed at or. The authors declare that they have no conflicts of interest or competing interests related to the manuscript. The opinions and assertions expressed herein are those of the author(s) and do not necessarily reflect the official policy or position of the Uniformed Services University or the Department of Defense.

## Abstract

Microglia, the resident immune cells of the central nervous system, have been subject to intense scrutiny recently because of their implicated role in the pathogenesis of Alzheimer’s disease, as well as many other neurological conditions. However, little is known about the diversity of microglial phenotypes, apart from the clear recognition that the M1-M2 distinction is an oversimplification. We used mass cytometry, a high dimensional analog of flow cytometry involving mass spectrometry-based analyses of single cells bound with metal-tagged antibodies, to characterize the diversity of microglia isolated from the brains of 3xTg-AD mice at 12, 18, and 24 months of age. Antibodies to microglial surface antigens CD11b, CD11c, CD14, CD18, CD24, CD35, CD38, CD45, CD51, CD64, CD68, CD86, CD124, CD172a, CD200R, CD206, CD274, CD301, CX3CR1, F4-80, MERTK, MHC class II, Siglec F, and TLR4 as well as intracellular antigens Arginase-1, IL10, IL12p40, iNOS, NADPH Oxidase, TGF-beta, and TNF-alpha were conjugated to unique metal isotopes, and used for mass cytometry. Phenograph-based clustering of the resulting high dimensional datasets identified 28 cell clusters of widely varying sizes, including clearly discrete phenotypes as well as pseudo-continuous/fractal phenotypic diversity. Monte Carlo simulations involving random reshuffling of the datasets revealed very different and much more uniform clustering patterns. Furthermore, control analyses of mouse peripheral blood mononuclear cells and brain microglia using the same markers revealed largely disjoint clusterings, indicating that these microglial clusters were not the result of peripheral blood contamination. Most of the microglial clusters were similar in samples from 3xTg-AD mice at all 3 ages, as well as from 12-month-old wild-type mice. The largest cluster expressed high levels of canonical microglial markers CD11b, CD11c, CD18, CD64, CD68, CD200R, CX3CR1, F4-80, NADPH oxidase, and TGF-beta but did not vary between experimental groups. However, 3 clusters did differ across experimental groups, one of which increased significantly with age in the 3xTg-AD mice as a fraction of the total microglial population. This cluster expressed high levels of CD18, CD38, CD45, CD86, and CD200R, but not other canonical markers. In summary, these mass cytometry-based analyses revealed previously uncharacterized microglial phenotypic heterogeneity in a mouse model of Alzheimer’s disease. The specific roles of the putative subclasses of microglia, their relevance to human disease, and many other questions remain to be addressed.

## Introduction

Originally identified by Pio del Rio Hortega around 1920, microglia are brain cells which perform functions similar to those of peripheral macrophages including phagocytosis of foreign materials and clearing debris.^1-3^ More recently it has been discovered that their adult phenotype is highly polymorphic, with changes directly related to the gut microbiota and many other stimuli.^1,4^ Microglia have furthermore been implicated in neurodegenerative diseases such as Alzheimer’s, and efforts are being made to determine how they might be altered to reduce the negative effects of these disorders.

Alzheimer’s disease is a rapidly growing threat and is currently the leading cause of dementia among the elderly. It is characterized by a loss of memory and other cognitive impairments, accompanied pathologically by the accumulation of tau aggregates, amyloid β (Aβ) plaques, and progressive brain atrophy.^5^ Microglial cells play a dual role in the presence of Aβ: they have been shown to produce potentially deleterious inflammatory factors, but also contribute to the clearance of Aβ through phagocytosis.^6,7^

The activation of microglia is the topic of intense study. A traditional view was that there were two activated microglial phenotypes: M1 (classically activated) and M2 (alternatively activated). The M1 classification was associated with neuroinflammatory effects, and M2 with neuroprotective effects. It is becoming apparent, however, that this view of microglial phenotypes is an oversimplification.^2^ At present, it is unknown exactly how many phenotypes of microglia exist or whether microglial activation can even be categorized into discrete phenotypes.

Mass cytometry is a newer technique that is being used to develop a deeper understanding of cells and their complex phenotypes.^8^ Flow cytometry is limited to modest numbers of parameters without introducing a large amount of spectral overlap. Mass cytometry allows the measurement of over 30 parameters on a single cell basis, with very little signal overlap. This method has been used to study many questions in the contexts of health, disease and pharmacologic intervention, ^8-12^ including the infiltrating blood-derived immune cell populations in the mouse brain, choroid and meninges.^13^

To explore the phenotypic complexity of microglia in the context of Alzheimer’s disease-like pathology, we used mass cytometry to examine the expression of 31 markers on microglial cells from wild type mice and Alzheimer’s disease model mice at 3 different ages.

This study was planned and executed without knowledge of the technically sophisticated mass cytometry studies of microglia recently published by Ajami et al.^14^ and Mrdjen et al.^15^

## Methods

### Animals

A total of 77 homozygous 3xTg-AD mice ^16^ (B6;129-*Psen1^tm1Mpm^* Tg(APPSwe,tauP301L)1Lfa/Mmjax, Cat#34830-JAX), were housed in the barrier facility at Washington University until the ages of 12, 18, and 24 months. A total of 25 wild-type mice (C57Bl/6J) at 12 months of age were purchased directly from Jackson Laboratory (Cat# 000664). Mice were acclimated to a 12 hour light-dark cycle, and food and water was provided ad libitum. Mice were housed at 3-5 animals per cage. Male and female mice were used in approximately equal numbers. All procedures involving animals were performed in accordance with the Washington University Animal Studies Committee Protocol # 20160030, Animal Welfare Assurance # A-3381-01.

### Perfusion and Brain Extraction

Groups of 4-5 animals were deeply anesthetized with isofluorane for 2 minutes. Mice were then transcardially perfused using ice cold 1x PBS (Diluted from 10x PBS, Leinco Technologies, MO) with 0.3% Heparin (Sagent Pharmaceuticals, IL). The animals were then decapitated and their brains removed. Olfactory bulbs and hindbrain were excised, and the remaining portion was immersed into 3 mL Serum-Free Media (Dulbecco’s Modified Eagle Medium (DMEM, Sigma-Aldrich, MO), 100 U/mL Penicillin-Streptomycin, 0.0045 mg/mL Glucose, and 0.01 M HEPES Buffer) at room temperature.

### Microglial Cell Isolation

Microglia were isolated using the Percoll gradient-based methods of Lee et al. ^17^ (**Figure 1A**). We chose to use this density-based method, rather than selection based binding to specific marker(s) such as CX3CR1 or CD11b, to reduce the likelihood of bias in favor of microglia which express higher levels of the specific marker(s). Immediately following perfusion and extraction, mouse brains were diced and placed in 3mL (per brain) of Dissociation Media (DMEM, 1x Hank’s Balanced Salt Solution (1xHBBS, Gibco, MO), 0.1 g/mL Papain (Sigma-Aldrich), 5 mg/mL Dispase II (Sigma-Aldrich), and 20 U/mL DNase I, Sigma-Aldrich). The Dissociation Media used equal volumes of 1x HBSS and DMEM, and DNase I was mixed with 1x HBSS, aliquoted, and added immediately before use. The tubes were then placed in a 37°C bath and mixed every 5 minutes by gentle inversion. 3 mL of Neutralization Media (DMEM, 100 U/mL Penicillin-Streptomycin (Gibco), 0.01M HEPES Buffer (Corning, NY), 0.0045 mg/mL Glucose (Sigma-Aldrich), and 10% v/v Fetal Bovine Serum (Gibco) per brain was added to stop enzyme activity, followed by centrifugation for 5 minutes at 23°C and 250 x *g*. Following aspiration of the supernatant, 3 mL of Serum-Free Media per brain was added to resuspend the remaining pellet, and the sample was spun down at the same settings. 3 mL of DMEM per brain was added and the sample was triturated using a glass Pasteur pipette which was fired to smooth the edges. Supernatant from the trituration was passed through a 70 μm cell strainer into a new tube and 3 mL of fresh DMEM was added. The process was repeated with a pipette fired to a reduced inner diameter than prior (approximately 0.75mm), and then a third time with a further reduction in inner diameter (approximately 0.5mm), each time running the supernatant through the cell strainer into a new tube to separate microglial cells from the tissue debris. The tube containing supernatant was then spun down at 250 x *g* and 23°C for 4 minutes. After another wash with DMEM, the sample was resuspended in 4 mL of 37% Stock Isotonic Percoll (90% Percoll (Sigma-Aldrich) by volume, 10% 10x HBSS by volume), diluted with 1x PBS, per brain. 4 mL of 70% Stock Isotonic Percoll per brain was underlaid (carefully deposited under the 37% layer) and 4 mL of 30% Stock Isotonic Percoll per brain was overlaid (carefully deposited over the 37% layer), with 2 mL of 1x HBSS per brain overlaid on top of that. The gradient was spun for 40 minutes at 18°C and 300 x *g*, allowing the cells to sediment in between the 30% and 37% layers. The interphase was then recovered using a transfer pipette. Approximately 5 mL were removed to guarantee that the entirety of the interphase was collected. The interphase fraction was transferred to a 15 mL conical tube and the remaining volume was filled with 1x HBSS and spun at 500 x *g* and 4°C for 10 minutes. The rinsed cells were resuspended and counted using a hemocytometer with trypan blue counter staining. Typical preparations yielded between 50,000, and 300,000 single living cells.

**Fig. 1.**
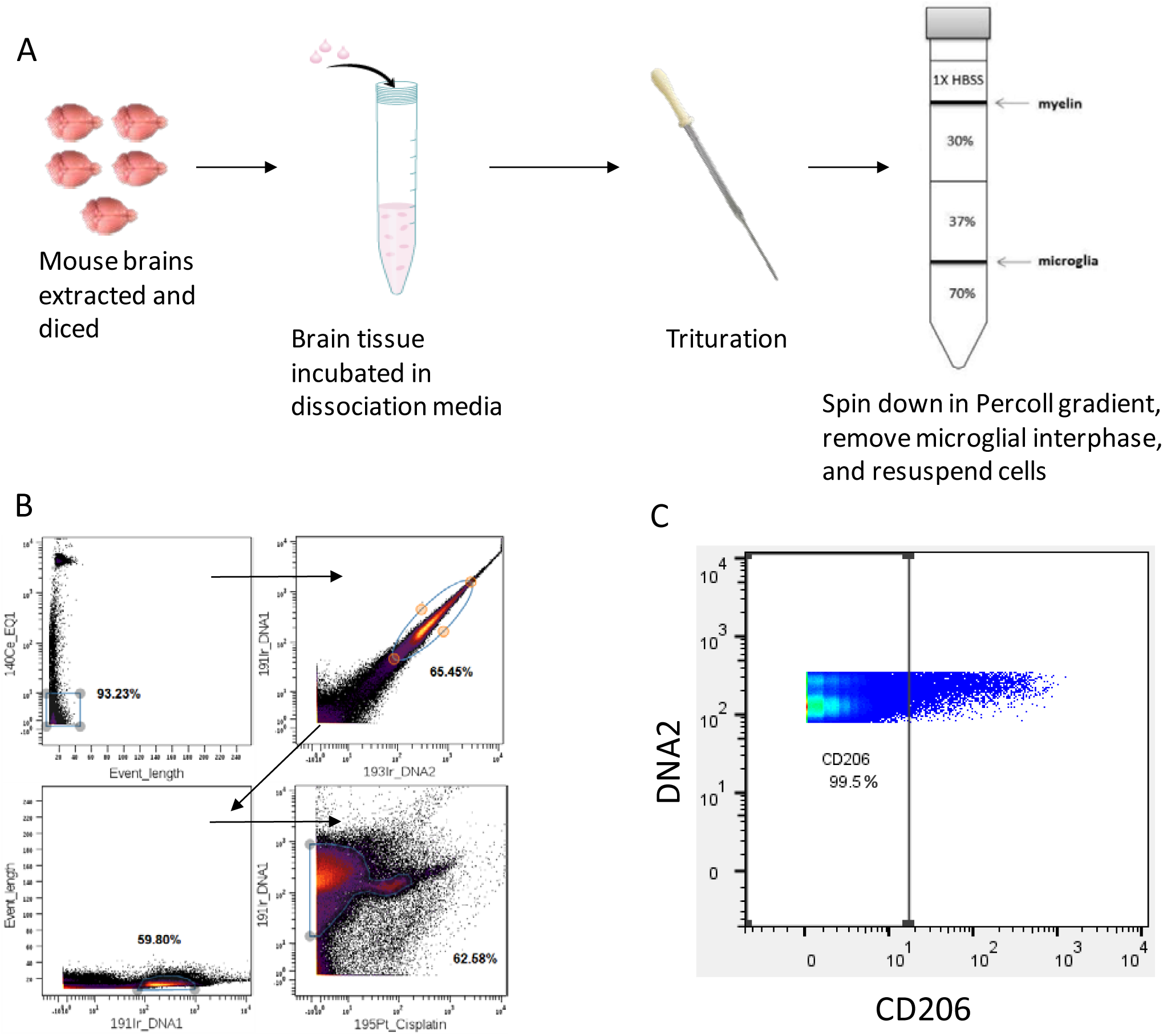
Illustration of Methods. **(A)** Microglial isolation protocol involving brain extraction, enzymatic dissociation, trituration, and Percoll gradient separation. (**B**) Gating for live, single cells. Events high in EQ1 indicate normalization beads, so those are gated out. Next, events with high DNA content based on Iridium DNA intercalator are gated in. Then single cells are gated in based on high DNA content and appropriate size (event length). Finally, live cells are gated in based on exclusion of cisplatin. (**C**) Gating out the 0.5% of the cells containing the highest expression of each marker (CD206 shown here) to improve data normalization. Data are from 12-month-old wild-type mice.

### Antibody Staining for Flow Cytometry to Validate Antibody Binding

Microglia from wild-type mice were resuspended in Blocking Buffer (1x PBS, 0.01M HEPES Buffer, and 1% v/v Normal Goat Serum (Vector Labs, CA)) and incubated on ice for 30 minutes in 1.5 mL microcentrifuge tubes (United Laboratory Plastics, MO). Staining Buffer (1x PBS, 1% v/v Normal Goat Serum, 0.01M HEPES Buffer, and Bovine Serum Albumin (Jackson Laboratories, CA)) was added and the cells were spun down. Tubes were spun for 30 seconds at room temperature, rotated 180°, then spun for another 30 seconds. The sample was resuspended in fresh Staining Buffer and aliquoted into separate samples for antibody incubation. The cells were equally divided into aliquots based on how many secondary antibodies were being used. (e.g. 10 secondary antibodies = 10 samples) Primary antibody was added and the samples were incubated on ice for 30 minutes. After another wash with Staining Buffer, Alexa Fluor® 488-conjugated secondary antibody, appropriate to the species of the primary antibody, was added at 0.125 to 0.5 μg/ml and the cells were incubated on ice for another 30 minutes. After three more washes with staining buffer were performed, the cells were resuspended in Staining Buffer. Flow cytometry was performed using a LSR Fortessa X-20 and analyses were performed after manual gating of viable single cells by forward scatter and side scatter criteria.

### Metal Conjugation

When available, antibodies were purchased from Fluidigm with metal conjugates (**Table 1**), For those that were not available, we utilized the Maxpar Antibody Labeling Kit according to the Fluidigm standard protocol (https://www.fluidigm.com/reagents/proteomics/201163a-maxpar-x8-antibody-labeling-kit--163dy--4rxn) to conjugate each antibody to a unique metal tag (**Table 1).**

**Table 1.** Antibodies used for mass cytometry.

| Antigen Name | Company | Catalog Number | Clone | Metal Tag | Antigen Location | Approx. Concentration in Cocktail* |
| --- | --- | --- | --- | --- | --- | --- |
| CD45 | Fluidigm | 3089005B | 30-F11 | 89Y | Surface | .25mg/mL |
| CD11c | Fluidigm | 3142003B | N418 | 142Nd | Surface | .5mg/mL |
| Siglec F | BD Biosciences | 552125 | E50-2440 | 143Nd | Surface | .5mg/mL |
| CD301 | Biolegend | 145702 | LOM-14 | 144Nd | Surface | .5mg/mL |
| TLR4 | Biolegend | 145402 | SA15-21 | 147Nd | Surface | .5mg/mL |
| CX3CR1 | Biolegend | 149002 | SA011F11 | 148Nd | Surface | .5mg/mL |
| CD24 | Fluidigm | 3150009B | M1/69 | 150Nd | Surface | .33mg/mL |
| CD64 | Fluidigm | 3151012B | X54-5/7.1 | 151Eu | Surface | .33mg/mL |
| CD274 | Fluidigm | 3153016B | 10F.9G2 | 153Eu | Surface | .5mg/mL |
| CD11b | Fluidigm | 3154006B | M1/70 | 154Sm | Surface | .05mg/mL |
| CD14 | Fluidigm | 3156009B | Sa14-2 | 156Gd | Surface | .5mg/mL |
| F4-80 | Fluidigm | 3159009B | BM8 | 159Tb | Surface | .5mg/mL |
| CD35 | BD Biosciences | 558768 | 8C12 | 160Gd | Surface | .5mg/mL |
| CD172a | BD Biosciences | 552371 | P84 | 163Dy | Surface | .125mg/mL |
| MERTK | Biolegend | 151502 | 2B10C42 | 165Ho | Surface | .5mg/mL |
| CD206 | Biolegend | 141702 | C068C2 | 167Er | Surface | .5mg/mL |
| CD124 | BD Biosciences | 551853 | mIL4R-M1 | 168Er | Surface | .5mg/mL |
| CD51 | Biolegend | 104102 | RMV-7 | 169Tm | Surface | .5mg/mL |
| CD68 | Biolegend | 137002 | FA-11 | 170Er | Surface | .5mg/mL |
| CD38 | Fluidigm | 3171007B | 90 | 171Yb | Surface | .25mg/mL |
| CD86 | Fluidigm | 3172016B | GL1 | 172Yb | Surface | .5mg/mL |
| MHC class II | Fluidigm | 3174003B | M5/114.15.2 | 174Yb | Surface | .018mg/mL |
| CD18 | Biolegend | 101402 | M18/2 | 175Lu | Surface | .25mg/mL |
| CD200R | Biolegend | 123902 | OX-110 | 176Yb | Surface | .5mg/mL |
| TNF-alpha | Fluidigm | 3141013B | MP6-XT22 | 141Pr | Intracellular | .5mg/mL |
| IL10 | Biolegend | 505029 | JES5-16E3 | 146Nd | Intracellular | .5mg/mL |
| IL12p40 | BD Biosciences | 554477 | C15.6 | 149Sm | Intracellular | .5mg/mL |
| NADPH Oxidase | BD Biosciences | 611415 | 53 | 152Sm | Intracellular | .5mg/mL |
| iNOS | Fluidigm | 3161011B | CXNFT | 161Dy | Intracellular | .5mg/mL |
| TGF-beta | BD Biosciences | 555052 | A75-2 | 164Dy | Intracellular | .5mg/mL |
| Arginase-1 | BD Biosciences | 610708 | 19 | 166Er | Intracellular | .05mg/mL |

\*concentrations of Fluidigm antibodies are not disclosed precisely.

### Preparation of Cisplatin

Cisplatin stock (100 mM aliquots) was thawed to room temperature, diluted 1:10 in PBS, and spun down at 12,000 x *g* for 5 minutes. This 10mM Cisplatin was aliquoted and frozen for future use. For use in experiments, 5μL of Cisplatin was added to 20 mL of CyPBS such that the Cisplatin was used at 2.5 μM.

### Cell Surface Marker Staining for Mass Cytometry

Cellular surface marker staining was performed as previously described.^18^ A cocktail of all antibodies to cell surface markers was prepared and maintained on ice prior to use. Each cellular sample was spun twice in CyFACS buffer, made from 1x PBS, 0.002 M EDTA, 1 mg/mL Bovine Serum Albumin, and 20 μg/mL Sodium Azide (Sigma-Aldrich), at 300 x *g* and 4°C for 5 minutes. The surface antibody cocktail was then added and the cells were incubated for one hour at 4°C. After the incubation, the cells were washed with CyPBS (10x PBS diluted to 1x) and spun at 500 x *g* and 4°C for 5 minutes. 2.5 μM cisplatin was added and mixed on a shaking rack for one minute. Following two additional washes with CyFACS using the same settings, the cells were incubated in 2% Paraformaldehyde (Electron Microscopy Sciences, PA) at 4°C overnight.

### Intracellular Marker Staining for Mass Cytometry

Intracellular antibody cocktail was prepared and stored on ice. The cells were washed twice using 1x Permeabilization Buffer (eBioscience). Chilled antibody cocktail was added to the cells and incubated at 4°C for one hour. After one more spin in CyFACS at 4°C and 800 x *g* for 5 minutes, the cells were incubated for 30 minutes at room temperature in 2% Paraformaldehyde with 500 μM Ir-Intercalator (Maxpar) diluted 1:2000 in CyFACS.

*concentrations of Fluidigm antibodies are not disclosed precisely.

## Analytical and Statistical Methods

### Gating and Normalization of Data

The initial gating was performed to separate out single, live cells. The order of gating was as follows: normalization beads were gated out, then a gate was placed around cells with high DNA content to exclude cellular debris and red blood cells, single cells were separated from clusters of cells based on “event length parameter” and finally, a live population was gated using cisplatin exclusion as a marker of viability (**Figure 1B**) according to standard methods.^18^ Once a live population was established, we gated out cells with expression levels in the top 0.5% for each marker (**Figure 1C**). This was performed because we found that outliers biased the normalization and can lead to less robust analytical results in Phenograph. We then normalized expression levels by the grand mean value for each marker across all valid datasets.

### Analysis of Mass Cytometry Data with Phenograph

We used the R package “cytofkit” for analyzing data with Phenograph. This package allowed us to use several FCS files at once, choose which markers would be used for clustering, and set various parameters such as the transformation, visualization, and clustering methods. We performed visualization with t-SNE (t-distributed Stochastic Neighbor Embedding) and clustering with Phenograph.^12^ The k-value, a parameter indicating the number of nearest neighbors assessed for each cell, was 75. Preliminary analyses indicated that smaller k-values led to larger numbers of clusters, and k-values larger than 75 did not affect the results. Our data had previously been transformed using hyperbolic arcsine with a cofactor of 5 prior to normalization, so we did not use any additional transformation here.^8^ We ran Phenograph on 5000 cells from each of 18 data sets (5 sets from 12 month 3xTG mice, 4 sets from 18 month 3xTG mice, 4 sets from 24 month 3xTG mice, and 5 sets from 12 month old WT mice). Each data set included microglia from 4-5 brains. Larger numbers of cells from each data set did not affect the results and substantially increased computational time. We also chose to remove 7 markers from the clustering pool; 6 (TNFα, IL10, TLR4, CD35, MERTK, CD124) were removed because their average expression did not differ between ages of 3xTg-AD mice, and the other marker (Arginase-1) was removed because there were irregularities involving very high binding during a subset of wild type mouse experiments. We ran each analysis several times using different randomly generated starting seeds, and found that the fundamental results were generally similar, though the shapes of the t-SNE plots differed.

### Monte Carlo Simulations

One set of data from 12-month-old 3xTG mice was used to run Monte Carlo simulations. Using the Microsoft Excel extension *Ablebits*, transformed expression levels for each marker were shuffled randomly, to remove any phenotypic relationships. This resulted in a table of mock cells with random expression levels but marginal distributions identical to the real data. This table was then converted into an FCS file and run through Phenograph to determine if it would find patterns where none should exist. This was done 5 times on the same set of data to produce 5 different random data sets, which in turn produced 5 different Phenograph plots.

## Results

We validated each of the antibodies used for the analyses by flow cytometry using microglia isolated from wild-type mice. Exemplars are shown in **Figure 2**.

**Figure 2.**
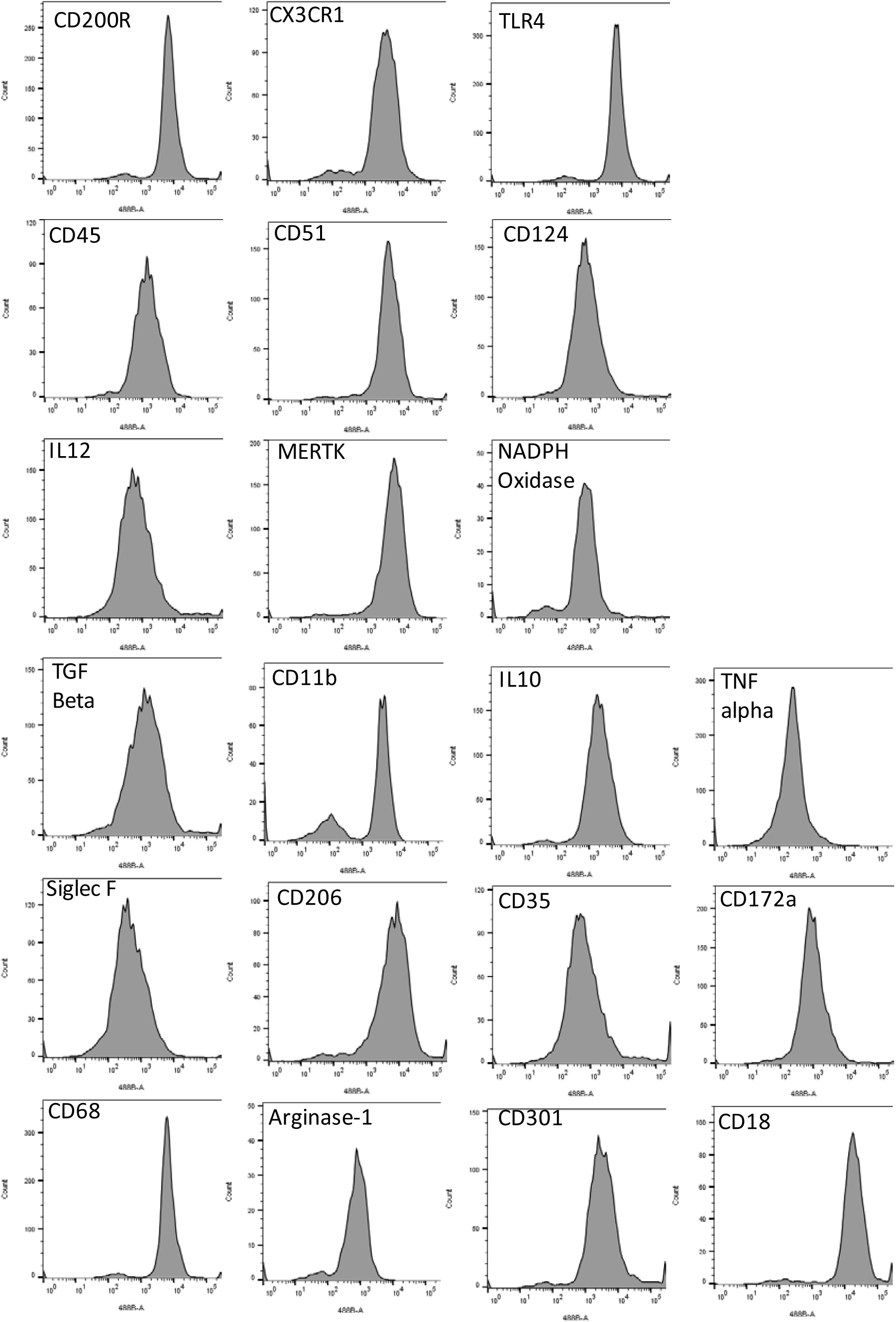
Exemplar Flow Cytometry Validation of the Antibodies Used in Mass Cytometry Studies. Secondary antibodies were conjugated to Alexa488. Microglia were isolated from wild-type mice. Unstained cells had fluorescence peaking under 2×10^2^.

### Microglial Marker Expression in 3xTg-AD Mice as a Function of Age

There was considerable variability in microglial marker expression as a function of age in 3xTg-AD mice, and differences between wild-type and 3xTg-AD mice at the same age (**Figure 3**). Notably, levels of CD14, CD18, CD38, CD68, CD86, CD200R, CD274, MHCII, and CX3CR1 all increased significantly with age. Levels of other canonical microglial markers including CD11c, F4-80, CD11b and the remainder of the markers did not change significantly as a function of age in 3xTg-AD mice. WT mice expressed modestly higher levels of several markers compared to 3xTg-AD mice at the same age (12 months) including CD38, F4-80, CD301 and CD124. However, the WT mice were not precisely strain matched to the 3xTg-AD mice, nor housed under identical circumstances, so these results should be interpreted cautiously. The units are arbitrary and do not directly reflect quantitative expression levels between different markers; they should only be interpreted for comparisons of expression level of the same marker between groups.

**Figure 3:**
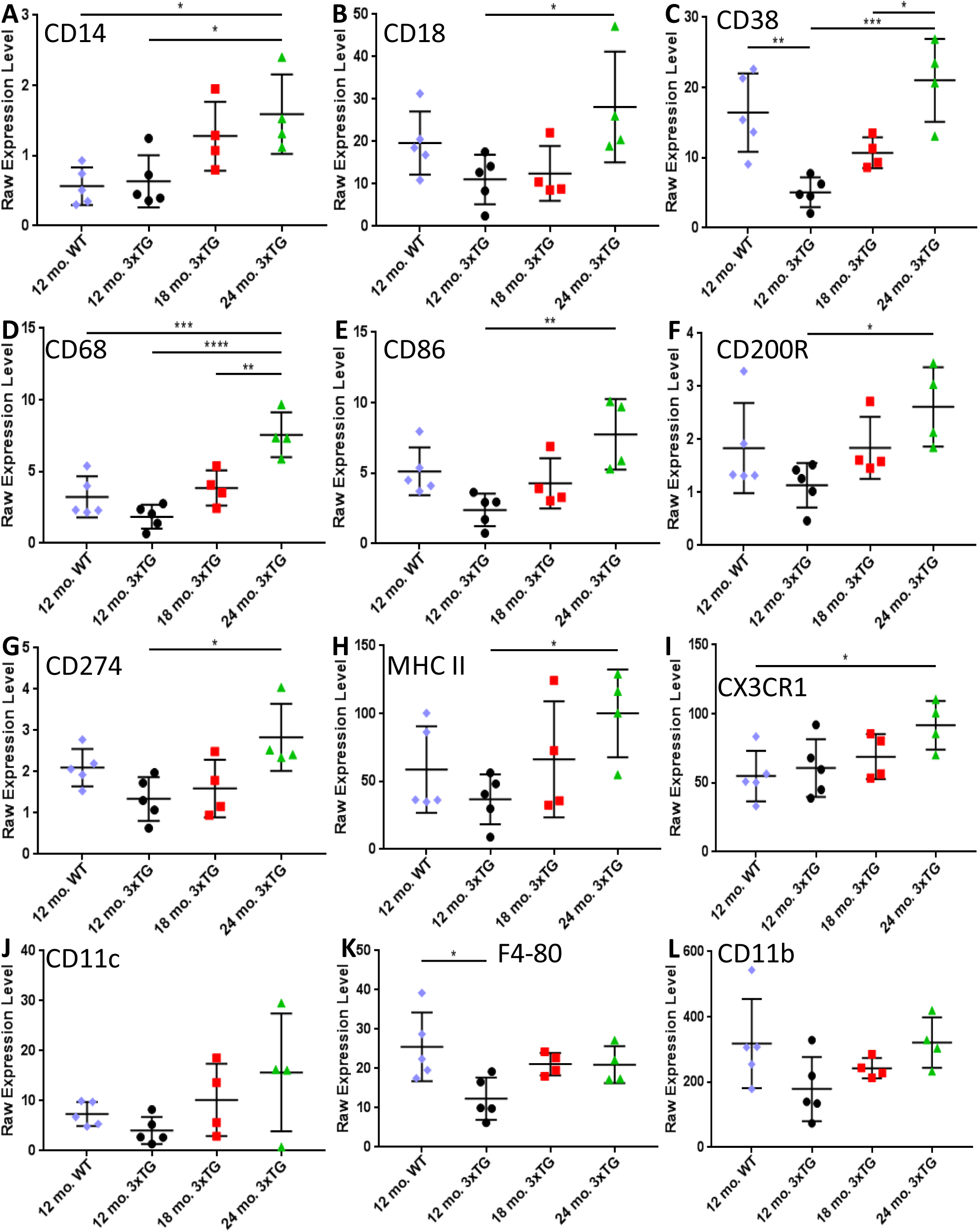
Mass Cytometry Microglial Marker Raw Expression Levels. (**A**) CD14: WT vs. 24 mo. 3xTG: *p=.0135, 12 mo. 3xTG vs. 24 mo. 3xTG: *p=.0213. (**B**) CD18 *p=.04. (**C**) CD38 *p=.0208, **p=.0048, ***p=.0004. (**D**) CD68 **p=.0052, ***p=.0009, ****p<.0001. (**E**) CD86 **p=.0027. (**F**) CD200R *p=.0250. (**G**) CD274 *p=.0141. (**H**) MHC II *p=.0435. (**I**) CX3CR1 *p=.0450. (**J**) CD11c, (**K**) F4-80 *p=.0185. (**L**) CD11b. All p-values indicate results of Tukey post-hoc comparisons after 1-way ANOVAs. The raw expression level are arbitrary units proportional to the number of ions detected and do not directly reflect quantitative expression levels between different markers. Each symbol represents a single experiment involving microglia pooled from 4-5 mouse brains.

### Microglial Clustering in 3xTg-AD Mice as a Function of Age

Phenograph-based clustering of microglia included both clearly discrete groups and nearly continuous / fractal phenotypic diversity (**Figure 4**). The algorithm divided the microglial cells into 28 clusters. There were clear divisions between many clusters (**Figure 4A**), indicating a dissimilarity in their expression profile (e.g. clusters 4, 25 and 28), but there were also clusters that were quite similar to their neighbors (e.g. clusters 6, 7, 10, 16 and 22), indicating a continuum of phenotypes. 5000 cells per experiment were used for computational tractability, but similar results were obtained for larger numbers of cells in individual experimental groups.

**Figure 4.**
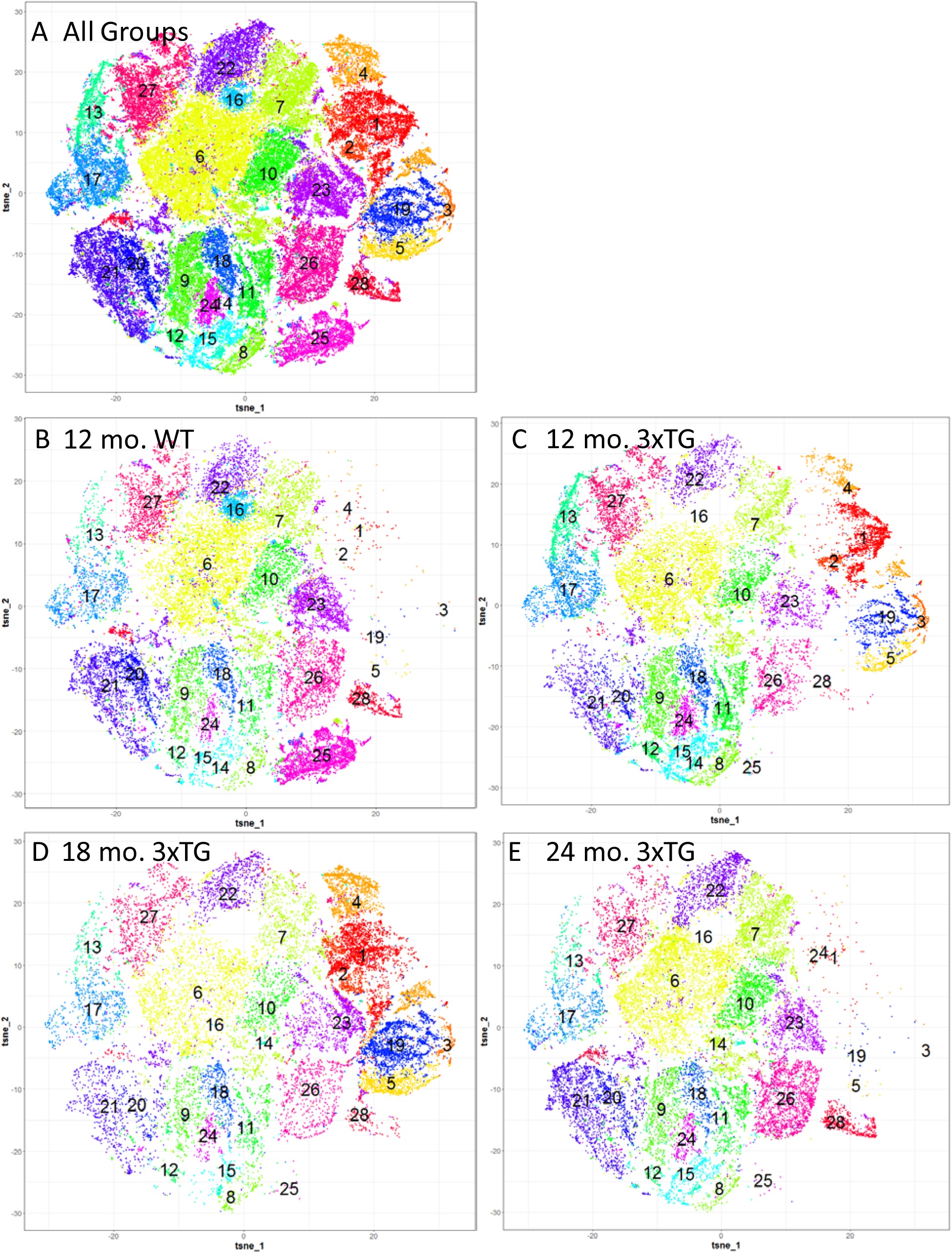
Phenograph Clustering of Microglia and Visualization Using t-SNE from Mass Cytometry Data. 5000 cells were randomly sampled from each experiment involving microglial isolated from 4-5 mouse brains. (**A**) All groups pooled with 28 clusters. (**B**) 12-month-old wild-type mice, 5 separate experiments pooled. (**C**) 12-month-old 3xTG mice, 5 separate experiments pooled. (**D**) 18-month-old 3xTG mice, 4 separate experiments pooled. (**E**) 24-month-old 3xTG mice, 4 separate experiments pooled. For 8 clusters (1, 2, 3, 4, 5, 16, 19, and 25), only a few experiments had a substantial fraction of microglial cells and results were inconsistent between experiments (see Figure 5). The other 20 clusters were considered valid.

Cluster 6, representing the single largest fraction of microglia, was notable for high expression of the canonical microglial markers CX3CR1, CD11b, F4-80, CD18, CD11c, CD64, TGF-beta, CD68, and CD200R. Clusters 7 and 10 were similar to cluster 6, though with the notable difference that cluster 7 had high Siglec F expression and cluster 10 had high CD301 expression.

Microglia from each of the 4 types of mice, including WT (**Figure 4B**), 12-month-old 3xTg-AD (**Figure 4C**), 18-month-old 3xTg-AD (**Figure 4D**), and 24-month-old 3xTg-AD (**Figure 4E**) had an overall similar clustering pattern, but there were also several notable differences between them. Specifically, there were 3 clusters where there were statistically significant quantitative differences between groups of mice (**Figure 5**). For cluster 15 (**Figure 5A,** 1-way ANOVA F_3, 14_=3.38, p=0.048), there was a significantly higher fraction of the population in 12-month-old 3xTg-AD mice compared with 18-month-old 3xTg-AD mice (Tukey post-hoc p=0.035). 12-18 months of age is a time period in which amyloid-beta and tau pathology are developing ^16,19^. However, the 24-month-old 3xTg-AD mice did not differ from the other groups. Cluster 15 was notable for high expression of CD200R. For cluster 20 (**Figure 5B,** 1-way ANOVA F3, 14=6.83, p=0.0046) There were significant differences between WT mice and 3xTg-AD mice at both 12 and 18 months of age (Tukey post-hoc p=0.015 and 0.008). However, again the 24-month-old 3xTg-AD mice did not differ from the other groups. Cluster 20 was notable for high expression of CD200R CD45, and CD274. For cluster 28 (**Figure 5C,** 1-way ANOVA F3, 14=7.3, p=0.0035), the fraction was highest in the WT mice (Tukey post-hoc p=0.018 vs. 12-month-old 3xTg), significantly lower in the 12-and 18-month-old 3xTg-AD mice, and significantly elevated again in the 24-month-old 3xTg-AD mice (Tukey post-hoc p=0.0063 and 0.046 vs. 12-month-old and 18-month-old 3xTg mice respectively). Cluster 28 was notable for high expression of CD86, CD45, CD38, CD200R, and CD18. Interestingly, canonical microglial markers such as CX3CR1, CD11b, F4-80 had low expression in clusters 15, 20 and 28.

**Figure 5.**
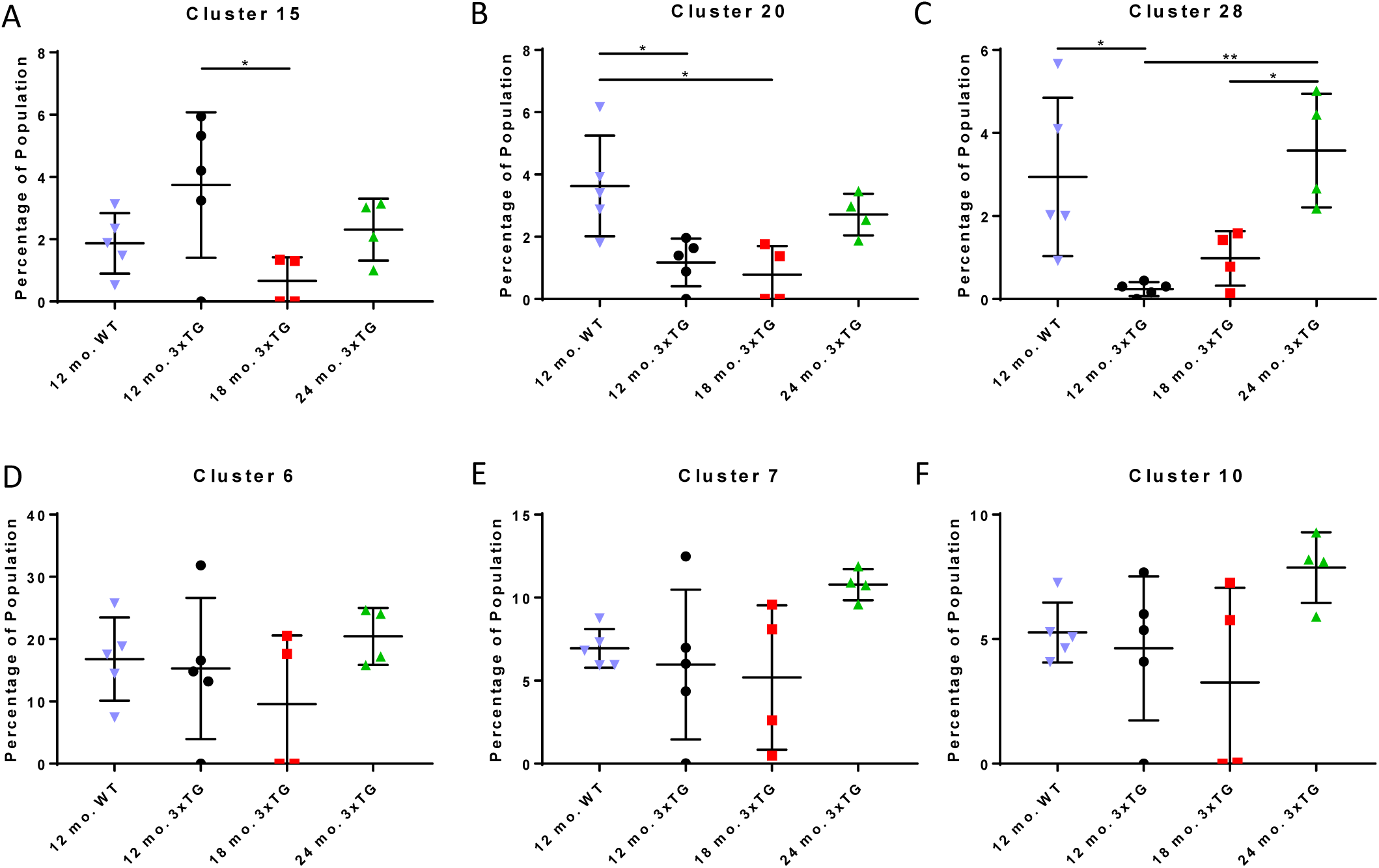
Quantitative Comparisons of the Percentage of Overall Microglia Represented by Phenograph-Derived Clusters. Percentages of each population organized by Phenograph cluster. (**A**) Cluster 15, notable for high CD200R expression, *p=.035. (**B**) Cluster 20, notable for high CD45 CD200R, and CD274 expression, *p=.0148, **p=.0079. (**C**) Cluster 28, notable for high CD18, CD38, CD45, CD86, and CD200R expression, *p=.0185 for 12 mo. WT vs. 12 mo. 3xTg, **p=.0063; *p=.0457 for 18 mo. 3xTg vs. 24 mo 3xTg. (**D**) Clusters 6, the largest cluster, notable for high expression of CD11b, CD11c, CD18, CD64, CD68, CD200R, CX3CR1, F4-80, NADPH oxidase, and TGF-beta. (**E-F**) Clusters 7 and 10, similar to cluster 6. All p-values indicate Tukey post-hoc comparisons after 1-way ANOVAs.

The other clusters of microglial cells identified by phenograph did not differ significantly between experimental groups. The clusters comprising the largest fraction of the microglia (cluster 6: ∼15-20% **Figure 5D**, clusters 7 & 10: ∼5-10% **Figure 5E-F**, and clusters 17, 21, 22, 23, 26, 27: ∼5% each, not shown) did not differ between experimental groups. For 8 clusters (1, 2, 3, 4, 5, 16, 19, and 25), only a few samples had a substantial fraction (>1%) of microglial cells. It is not clear why these samples differed from other identically handled samples from mice of the same genotype. Each sample was derived from 4-5 individual mice, so this variability is not likely to result from genetic mutations. However, often the mice in each sample came from the same cage, so infection or other immunological stimuli could have played a role. This will require future investigation outside the scope of the current report. Thus, we considered the other 20 clusters to be valid.

### Peripheral Blood Mononuclear Cells vs. Brain Microglia

As a control, we took samples of peripheral blood mononuclear cells (PBMCs) and microglia from our WT mice to compare the two. Our goal was to determine whether the cells isolated in our brain preparation are phenotypically similar to blood cells, either because of *in vivo* infiltration or *post-*mortem contamination. 10,000 cells were subsampled from each of 9 datasets (5 from microglia, 4 from PBMCs) and clustered using Phenograph. We found that the PBMCs and microglia occupied mostly different clusters with minimal overlap (**Figure 6**). This demonstrated that the microglial cells analyzed were not likely to have been substantially intermingled with PBMCs and that these two cell populations are phenotypically distinct.

**Figure. 6.**
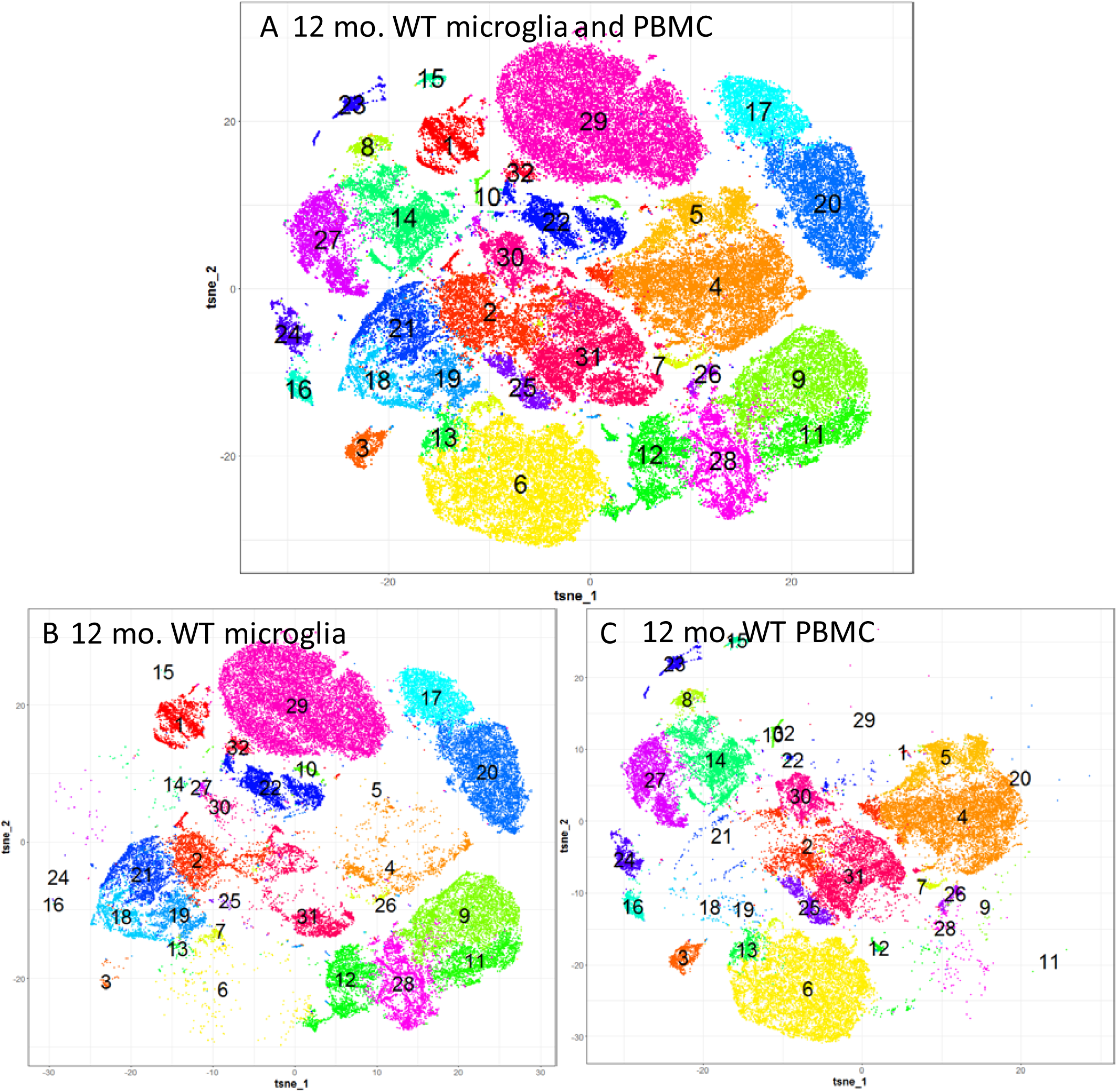
Microglia vs. Peripheral Blood Cells Compared by Mass Cytometry. Microglia and peripheral blood mononuclear cells (PBMCs) were isolated from the same wild-type mice. 10,000 cells were randomly sampled from 9 files (5 from wild-type microglia, 4 from wild-type PBMCs) (**A**) Both microglia and PBMCs together. (**B**) Microglial cells only. (**C**) PBMCs only.

### Phenograph Analyses of Data from Monte Carlo Simulations

As an additional control, we performed analyses of mock datasets derived from Monte Carlo simulations with the same marginal distributions as data from 12-month-old 3xTg-AD mice (**Figure 7**). Phenograph was able to create clusters based on random phenotypic similarity, but the shape and size of the clusters from the simulated data was very uniform (**Figure 7B-F**), in contrast to the clusters derived from the real microglial data (**Figure 7A**). The Monte Carlo-derived clusters also tended to have a uniform distance between them, unlike real microglial data sets which were characterized by tight groups of clusters and other substantially more distant clusters in the t-SNE plots. These results indicate that the clustering observed in the real microglial data sets was not likely to have been an artifact due to clustering by the Phenograph algorithm.

**Figure 7.**
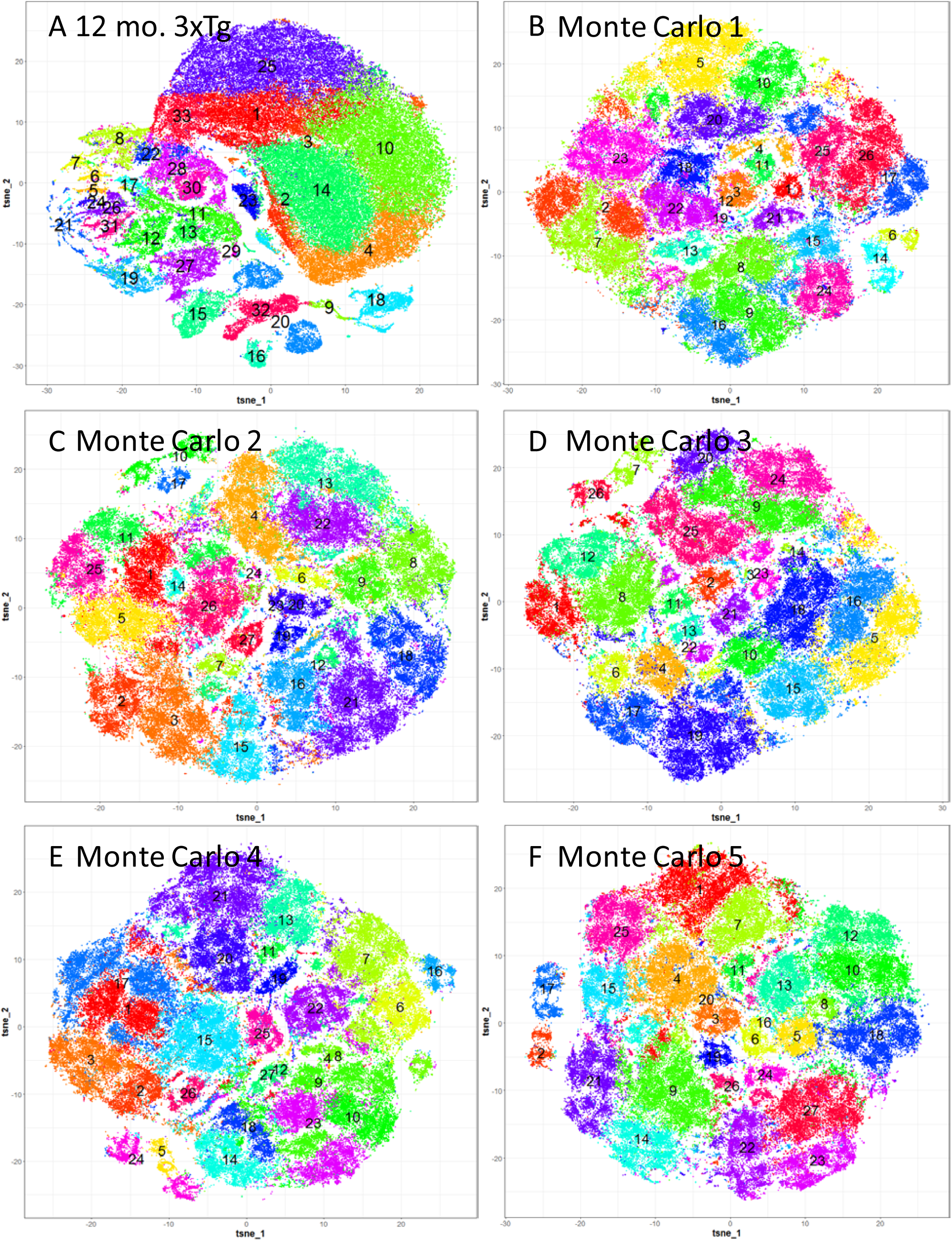
Phenograph Clustering of Mass Cytometry Data from 12-month-old 3xTG Mouse Microglia vs. Monte Carlo Simulations. (**A**) Phenograph clustering and t-SNE visualization of experimental data from 12-month-3xTG mice. (**B-F**) Monte Carlo simulations. The experimental data from panel A were randomized by randomly shuffling expression values for each marker. This was performed five separate times, and the results were analyzed using methods identical to those used for the experimental data. The experimental data with intrinsic correlations has a higher degree of clustering than the Monte Carlo simulations.

### Gradients of Expression Levels for Microglial Markers

In addition to the discrete clusters and sub-clusters of microglia found using Phenograph-based analyses, we also observed gradients of expression levels for several of the microglial markers based on the original t-SNE analyses. For example, within apparent clusters, CD11b and MHC II expression formed sharp gradients, which may differ as a function of age in 3xTg mouse microglia (**Figure 8**). We have not yet quantitatively analyzed the gradients as a function of experimental group, but report this as an observation warranting additional assessment.

**Figure 8.**
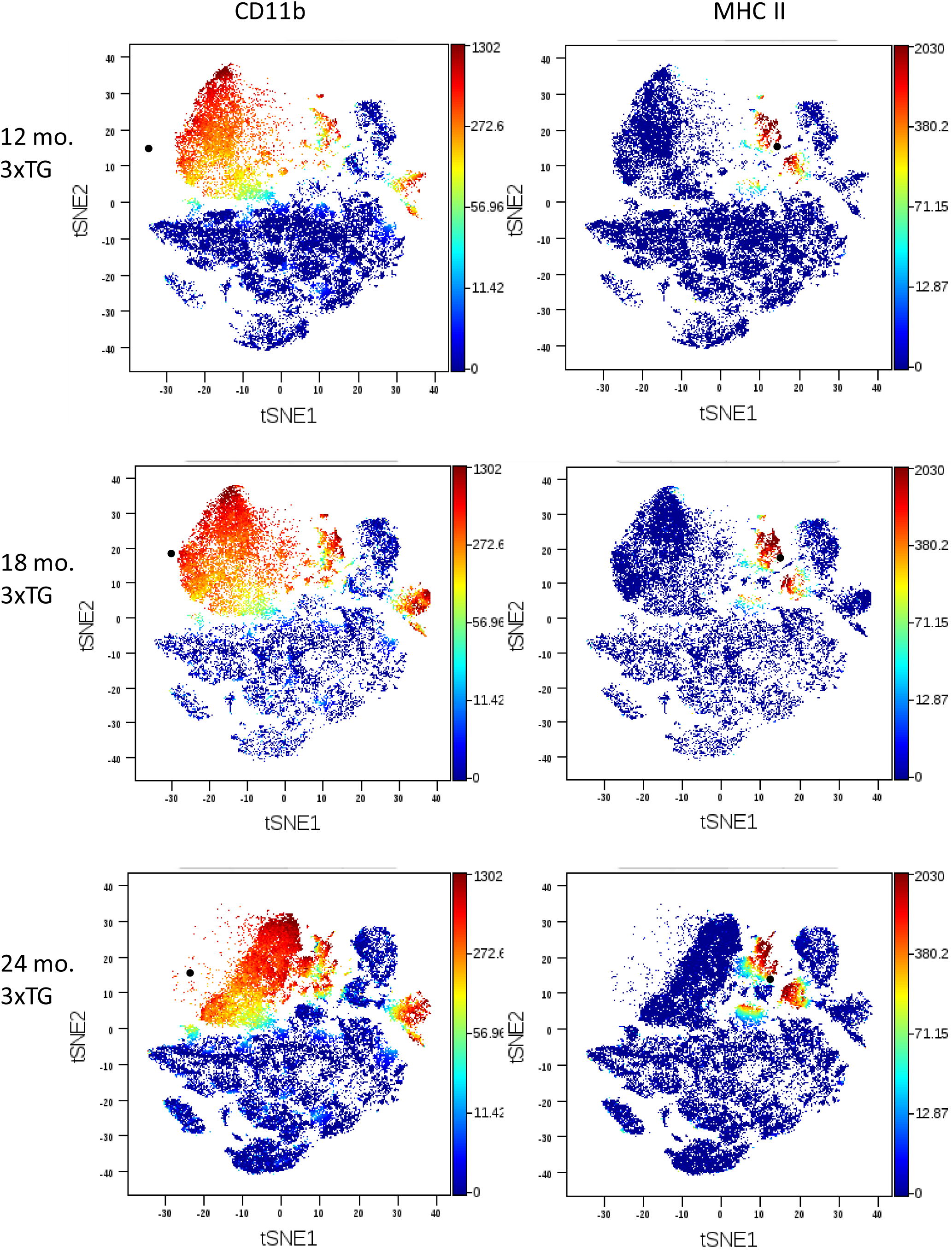
Gradients of Expression of CD11b and MHC II Based on Single Cell Mass Cytometry Data. Color scale represents raw expression levels transformed using hyperbolic arcsine with a cofactor of 5. Data from single experiments from 12-, 18- and 24-month-old 3xTG mouse microglia, with 4-5 mouse brains per experiment.

## DISCUSSION

We characterized mouse brain microglial phenotypes using mass cytometry, a high dimensional, antibody-based method. The pattern of expression of these markers was clearly more complex than can be explained by the M1-M2 dichotomy prevalent in the literature. The phenotypes can be characterized as a superposition of discrete populations and gradients of expression. We identified 28 discrete populations with characteristic markers expressed in each population, 20 of which appeared to be valid. The discrete populations in some cases also involved a hierarchical nesting of sub-populations expressing many but not all of the markers that characterized the populations as a whole. The number of clusters produced by the phenograph algorithm is somewhat arbitrary, and it is possible that there is no ‘correct’ number of clusters due to fractal sub-clustering. In addition, expression of some of the most common microglial markers such as CD11b, CD68, CX3CR1 and F4-80 appeared as gradients ranging from low to high across the populations, with some microglial cells negative for these markers. It is clear that microglia cannot be considered a unitary cell population, nor characterized by a simple dichotomy of phenotypic states. These results should be interpreted as a complement to gene expression studies ^20-23^, which while more unbiased and able to assess a broader range of genes, do not offer the opportunity to directly assess protein expression at a single cell level. Our study used a substantially different set of mass cytometry markers than the recently published, elegant studies in other mouse models of neurological disorders^14,15^. Thus, the results cannot be directly compared. Taken together, our results and the intriguing recently reported results in experimental autoimmune encephalomyelitis,^14,15^ Huntingtin transgenic mice,^14^ the mutant superoxide dismutase-1 transgenic mouse model of Amyotrophic Lateral Sclerosis,^14^ aged C57Bl6 mice,^15^ and the APP-PS1 mouse model of Alzheimer’s disease,^15^ indicate the substantial value of the mass cytometry approach to analysis of microglia.

While most of the discrete populations were similar in relative abundance between 3xTg-AD mice of different ages, a few did differ significantly with age. Unexpectedly, CD200R turned out to be widely expressed on the microglial clusters that varied as function of age in 3xTg-AD mice, even though these clusters did not express high levels of other canonical microglial markers. It should be noted again that the cells analyzed were based on Percoll gradient separation, may not all have been traditionally defined microglia. Nonetheless, further investigation into the role of CD200-CD200R signaling in the context of aging and AD-related pathology will be warranted. CD200-CD200R interaction may represent a ‘stop signal’ by which neurons and other brain cells prevent excessing microglial or other immune cell activation along pathways that lead to phagocytosis of surrounding material. ^24^ Changes in relative abundance of cell populations expressing high levels of CD200R could hypothetically affect the competence of microglia to engulf plaques, or aberrantly prune synapses. Furthermore, inescapable tail shock stress has been reported to cause downregulation of CD200R in hippocampus and amygdala in rats,^25^ though whether this contributed to the effects of stress on the development of AD pathology ^26-28^ remains to be determined. CD200R was by no means a universal marker for all microglial subsets in that its expression was low in clusters 1-5 and 19. Furthermore, expression of CD200R is not unique to microglia. Granulocytes, T cells, mast cells, and activated basophils have also been reported to express CD200R.^29^ Thus, CD200R should not be considered a specific ‘pan-microglia’ marker.

These results are not without limitations, some of which are elaborated here:

1. The number of clusters found using the Phenograph algorithm is somewhat arbitrary, and depends on the clustering parameter k. With other values of k, we did not observe major differences in the results, but we recognize that other groupings including fractal and branched sub-clustering organizational strategies could also be employed.
2. The antibodies selected were based on a combination of literature review and feasibility of binding under a uniform set of conditions for all antibodies; the use of additional antibodies may reveal even greater diversity, as could single cell expression profiling. Thus, even the tremendous complexity of expression patterns and diversity of microglial subpopulations revealed here may represent just the tip of the iceberg. For example, several groups have recently characterized individual microglia using single cell RNA sequencing and revealed additional diversity. ^20,23 22^ Regrettably, at the time the antibody panel was finalized, high quality TREM2 antibodies were not available to us and we did not include antibodies to TREM2 in the panel. TREM2 plays an important role in microglial function in the setting of AD ^30,31^, and future investigations should directly assess TREM2 expression across the spectrum of microglial phenotypes.
3. We cannot be certain that the microglial expression did not change during the perfusion, brain extraction, isolation and staining processes, which occurred over 36 hours even though all steps were performed on ice or at 4°C; orthogonal *in situ* methods will be required for validation.
4. We used a moderately restrictive set of gating parameters to strike a balance between excluding non-microglial cells and including sufficient cells for robust analyses; alternative gating parameter choices could affect these results.
5. We assessed relatively modest numbers of cells, and there was likely to have been substantial cell loss during the preparation. Thus, the extent to which the isolated and characterized cells are representative of the whole brain population of microglia is unknown. Alternative approaches such as Tmem119-based isolation should be considered in the future.^32^
6. Because yield of microglia varied from preparation to preparation, there was no way to assess the absolute numbers of microglia based on these data; all analyses were based on relative quantification. Alternative *in situ* methods with stereological quantification will be required to assess absolute numbers of specific microglial populations.
7. We have not evaluated microglial diversity in wild-type mice as a function of age, and the 12-month-old wild-type mice were used were not exactly strain matched to the 3xTg-AD mice. Characterization of additional lines of AD model mice with littermate controls would be an appropriate future direction.
8. We have not directly verified the specificity of the antibodies used, however, multiple markers were used to characterize each cluster, which should make the results relatively robust even if one or a few antibodies lack complete specificity.
9. While the preparation methods were designed to isolate microglia from other brain cells, we did not directly assess for non-microglial brain cells using markers of other lineages (e.g. GFAP, NeuN)
10. We have not performed perfusion controls as in Korin et al.^13^, though the results of our direct comparison between PBMCs and microglia in the same mice strongly suggests that our perfusions were largely adequate and that few PBMC were present in our microglial preparations.

Thus, clearly many additional experiments will be required to definitively assess microglial phenotypic diversity.

This initial report raises the possibility of addressing many additional scientific questions. Some of the questions include the following: 1) What are the functional phenotypes of these microglial subsets? Cell sorting followed by *in vitro* experiments could begin to address this question, as could transgenic animals expressing combinations of fluorescent reporters (e.g. “brainbow”^33^). 2) Do the subsets of microglia interconvert under physiological and pathophysiological conditions? Cell fate labeling *in vivo* followed by observations over time after various stimuli could be a relevant approach. 3) How does this diversity of microglia arise, i.e. are the subsets genetically defined or do they result from responses to environmental signals? 4) How do microglia of the various subclasses identified here interact with AD pathology ^34 35 36^ and which if any are responsible for aberrant synaptic pruning ^37^ *In situ* multi-label methods addressing these questions will be an important future direction. 5) How does the microglial phenotypic complexity in mice relate to that in the human central nervous system? Mass cytometry applied to microglia isolated from surgically resected tissue ^38^, or *in situ* scanning mass cytometry in tissue slices ^39^ could be used to address this question once suitable antibodies for human microglial antigens have been identified and validated. Interestingly, gene expression profiling of human brain tissue from patients with neurodegenerative disorders has revealed patterns not apparent in some animal models ^20^, though the extent to which these gene expression patterns drive microglial phenotypic diversity remains to be determined. 6) What really is a microglial cell? If microglia are defined by motility and surveillance function, then *in vivo* multiphoton microscopy using mice expressing fluorescent reporters tagging various microglial populations could be an appropriate approach for future inquiry.

Clearly, the current report raises more questions about microglia than it answers, but it provides a framework for many deep physiological and pathophysiological investigations. While the univariate results involving single markers could have been obtained using flow cytometry or comparable traditional methods, the potentially more interesting clustering results could not have been obtained without the use of mass cytometry or another high dimensional method.

Conclusion: Microglial diversity at least in mice appears far broader than traditionally appreciated. The diversity can be characterized as a mixture of discrete subpopulations and continuous gradients. Mass cytometry may represent an attractive tool for characterizing this diversity, and should be applicable to many other developmental and pathophysiological settings where microglia are believed to play important roles. It is hoped that new insights based on a deep understanding of microglial diversity will allow targeted therapeutic interventions to boost beneficial functions of microglia and dampen harmful microglial-mediated processes in disorders of the nervous system.

## Acknowledgements

The authors would like to thank the F-Prime foundation for financial support, Dr. Florent Ginhoux for early advice on this project, Dr. David Holtzman and his laboratory staff for use of their flow cytometer, Drs. Terrence Kummer and Andrew Sauerbeck for critical comments, and the Brody, Kummer and Friess lab members for stimulating discussions.

## Author Contributions

EMF performed the majority of the experiments, analyzed the majority of the data, and wrote the first draft of the manuscript.

TJE assisted with cell isolation and flow cytometry experiments and provided troubleshooting advice.

OM assisted with mass cytometry experiments and provided troubleshooting advice.

STO provided expertise regarding design and execution of mass cytometry experiments.

DLB conceived the project, obtained funding, supervised the project, provided troubleshooting advice, analyzed data, and wrote the final version of the manuscript.

All authors reviewed the manuscript.

